# MASSpy: Building, simulating, and visualizing dynamic biological models in Python using mass action kinetics

**DOI:** 10.1101/2020.07.31.230334

**Authors:** Zachary B. Haiman, Daniel C. Zielinski, Yuko Koike, James T. Yurkovich, Bernhard O. Palsson

## Abstract

Mathematical models of metabolic networks utilize simulation to study system-level mechanisms and functions. Various approaches have been used to model the steady state behavior of metabolic networks using genome-scale reconstructions, but formulating dynamic models from such reconstructions continues to be a key challenge. Here, we present the Mass Action Stoichiometric Simulation Python (MASSpy) package, an open-source computational framework for dynamic modeling of metabolism. MASSpy utilizes mass action kinetics and detailed chemical mechanisms to build dynamic models of complex biological processes. MASSpy adds dynamic modeling tools to the COnstraint-Based Reconstruction and Analysis Python (COBRApy) package to provide an unified framework for constraint-based and kinetic modeling of metabolic networks. MASSpy supports high-performance dynamic simulation through its implementation of libRoadRunner; the Systems Biology Markup Language (SBML) simulation engine. Three case studies demonstrate how to use MASSpy: 1) to simulate dynamics of detailed mechanisms of enzyme regulation; 2) to generate an ensemble of kinetic models using Monte Carlo sampling to approximate missing numerical values of parameters and to quantify uncertainty, and 3) to overcome issues that arise when integrating experimental data with the computation of functional states of detailed biological mechanisms. MASSpy represents a powerful tool to address challenge that arise in dynamic modeling of metabolic networks, both at a small and large scale.

**Author Summary:** Genome-scale reconstructions of metabolism appeared shortly after the first genome sequences became available. Constraint-based models are widely used to compute steady state properties of such reconstructions, but the attainment of dynamic models has remained elusive. We thus developed the MASSpy software package, a framework that enables the construction, simulation, and visualization of dynamic metabolic models. MASSpy is based on the mass action kinetics for each elementary step in an enzymatic reaction mechanism. MASSpy seamlessly unites existing software packages within its framework to provide the user with various modeling tools in one package. MASSpy integrates community standards to facilitate the exchange of models, giving modelers the freedom to use the software for different aspects of their own modeling workflows. Furthermore, MASSpy contains methods for generating and simulating ensembles of models, and for explicitly accounting for biological uncertainty. MASSpy has already demonstrated success in a classroom setting. We anticipate that the suite of modeling tools incorporated into MASSpy will enhance the ability of the modeling community to construct and interrogate complex dynamic models of metabolism.

## Introduction

The availability of genome sequences and omic data sets has led to significant advances in metabolic modeling at the genome scale, resulting in the rapid expansion of available genome-scale metabolic reconstructions [1]. COnstraint-Based Reconstruction and Analysis (COBRA) methods [2] have been shown to be a scalable framework that is invaluable for the contextualization and analysis of multi-omic data, as well as for understanding, predicting, and engineering metabolism [3–12]. While several methods have been developed that allow COBRA models to integrate certain data types to model long timescale dynamics [13–15], COBRA models are inherently limited by the flux-balance assumption.

Kinetic modeling methods use detailed mechanistic information to model dynamic states of a network [16]. The inclusion of multiple detailed enzymatic mechanisms presents challenges in formulating and parameterizing stable large-scale kinetic models. Further, additional issues arise when integrating incomplete experimental data into metabolic reconstructions, necessitating the need for approximation methods to gap fill missing values that satisfy the thermodynamic constraints imposed by the system [17, 18].

Various efforts have been made to bridge the gap between constraint-based and kinetic modeling methods in order to address the challenges associated with dynamic modeling [17–20]. One such methodology is the Mass Action Stoichiometric Simulation (MASS) approach, in which mass action kinetics are used to construct condition-specific dynamic models [20–23]. The MASS modeling approach provides an algorithmic, data-driven workflow for generating *in vivo* kinetic models in a scalable fashion [24]. The MASS methodology can be used in tandem with COBRA methods for both steady-state and dynamic analyses of a metabolic reconstruction in a single workflow. MASS models can incorporate the stoichiometric description of enzyme kinetic mechanisms and have been used to explicitly compute fractional states of enzymes, providing insight into regulation mechanisms at a network-level [21]. The MASS modeling framework has been implemented in the MASS Toolbox [25], but is limited by its reliance on a commercial software platform (Mathematica).

Here, we detail the Mass Action Stoichiometric Simulation Python (MASSpy) package, a versatile computational framework for dynamic modeling of metabolism. MASSpy expands the modeling framework of the COnstraint-Based Reconstruction and Analysis Python (COBRApy) [26] package by integrating dynamic simulation and analysis tools to facilitate dynamic modeling. Further, MASSpy contains various algorithms designed to address and overcome the issues that arise when incorporating experimental data and biological variation into dynamic models with detailed mechanistic information. By addressing the issues associated with integrating physiological measurements and biological mechanisms in dynamic modeling approaches, we anticipate that MASSpy will become a powerful modeling tool for modeling dynamic behavior in metabolic networks.

## Design and implementation

### Developing in Python

The MASSpy software package (S1 File) is written entirely in Python 3, an interpreted object-oriented high-level programming language with a clean syntax that has become widely adopted in the scientific community due to its unique features (e.g., a flexible interface to compiled languages such as C++ [27]). The open-source nature of Python avoids the inherent limitations associated with costly commercial software [28]. Consequently, developing in Python provides access to a growing variety of open-source scientific software libraries [29, 30], several of which are integrated into the MASSpy package and utilized for various purposes (Table 1).

**Table 1.** Overview of external dependencies and their relevance to MASSpy functionality.

| Package | Version | MASSpy Relevance | Reference |
| --- | --- | --- | --- |
| COBRApy | 0.15.0 | Reconstruction and simulation of genome-scale flux states | [26] |
| Escher | 1.7.2 | Visualization of pathway and node maps | [31] |
| libRoadRunner* | 1.5.0 1 | Dynamic simulation and steady state determination through NLEQ methods | [32] |
| libSBML* | 5.18.0 1 | A Python interface for reading and writing models in SBML | [33] |
| Matplotlib* | 3.2.0 | Visualization of simulation results | [34] |
| Numpy | 1.13.0 | Fundamental package for numerical computation in Python. Provides efficient array/matrix data types and operations. | [35] |
| OptLang | 1.4.2 | Formulation of optimization problems using symbolic expressions and native Python algebra syntax. Provides a common interface for various optimization solver backends. | [36] |
| Pandas | 0.17.0 | High-performance data structures and analysis tools for data science | [37] |
| SciPy* | 1.2.0 | A collection of scientific algorithms. Primarily used for interpolation of dynamic simulation results and linear algebra operations. | [38] |
| SymPy | 1.0.0 | Generation and manipulation of symbolic mathematical expressions, including ordinary differential equations, rate laws, and optimization problems. | [39] |
| swiglpk | 1.4.3 | A Python interface to the GNU Linear Programming Kit used for optimization. Utilized by OptLang to provide LP and MILP support. | [40] |
| cplex** | 12.8.8.0 | A Python interface to the CPLEX Optimizer used for optimization. Utilized by OptLang to provide LP, MILP, and QP support. | IBM,<br>Armonk, NY |
| gurobi** | 5.0.2 | A Python interface to the Gurobi Optimizer used for optimization. Utilized by OptLang to provide LP, MILP, and QP support. | Gurobi<br>Optimization,<br>Houston, TX |

\* Additional packages required to enable all MASSpy features; all other packages are strict or optional dependencies of COBRApy.
\*\* Commercial optimization solvers with Python APIs with free academic licenses. Abbreviations: Linear Programming (LP); Mixed Integer Linear Programming (MILP); Quadratic Programming (QP); Systems Biology Markup Language (SBML);

### Building on the COBRApy framework

To facilitate the integration of constraint-based and dynamic modeling frameworks, MASSpy utilizes the COBRApy package [26] as a foundation to build upon and extend in order to support dynamic simulation and analysis capabilities. MASSpy derives several benefits from building on the COBRApy framework, including exploiting the direct inclusion of various COBRA methods already implemented in Python. The inclusion of COBRA methods is made simple using Python inheritance behavior; the three core COBRApy classes (Metabolite, Reaction, and Model) serve as the base classes for three core MASSpy classes (MassMetabolite, MassReaction, and MassModel) as described in the MASSpy documentation (https://masspy.readthedocs.io/ and S2 File). Consequently, all methods for COBRApy objects readily accept the analogous MASSpy objects as valid input, preserving the commands and conventions familiar to current COBRApy users. COBRApy is a popular software platform preferred by many in the COBRA community [41]; therefore, preserving COBRApy conventions aids in the adoption of MASSpy among those users. Inheriting from the COBRApy classes additionally allows for easy conversion between COBRApy and MASSpy objects without loss of relevant biochemical and numerical information. These two features of Python inheritance are critical in maintaining functionality for COBRApy implementations of various flux-balance analysis (FBA) algorithms in MASSpy.

### Adding dynamic simulation capabilities

The creation and simulation of dynamic models requires deriving a set of ordinary differential equations (ODEs) from the stoichiometry of a reconstructed network and assigning kinetic rate laws to each reaction in the network [17]. MASSpy utilizes SymPy [39] to represent reaction rates and differential equations as symbolic expressions. All MassReaction objects automatically generate their own rate laws using mass action kinetics, unless a suitable rate law is available from literature and assigned to the reaction. All MassMetabolite objects generate their associated differential equation by combining the rates of reactions in which they participate and contain the initial conditions necessary to solve the system of ODEs.

To solve the system of ODEs, the MASSpy Simulation class employs libRoadRunner [32], a high-performance Systems Biology Markup Language (SBML) [42] simulation engine that is capable of supporting most SBML Level 3 specifications. The libRoadRunner utilizes a Just-In-Time (JIT) compiler with an LLVM JIT compiler framework to compile SBML-specified models into machine code, making the libRoadRunner simulation engine appropriate for solving large models effectively. Although libRoadRunner has a large suite of capabilities, it is currently used for two purposes in MASSpy: the steady-state determination via NLEQ1 and NLEQ2 global newton methods [43], and dynamic simulation via integration of ODEs through deterministic integrators, including CVODE solver from the Sundials suite [44]. Because libRoadRunner requires models to be in SBML format, the Simulation object exports models into SBML format before compiling them into machine code via libRoadRunner.

### Model import, export, and network visualization

MASSpy utilizes two primary formats for the import and export of models: SBML format and JavaScript Object Notation (JSON). MASSpy currently supports SBML L3 core specifications [45] along with the FBC [46] and Groups [47] packages, providing support for both constraint-based and dynamic modeling formats. Although SBML is necessary to utilize libRoadRunner, there are a number of additional benefits obtained by supporting SBML. In addition to being a standard format among the general systems biology community [42], SBML is a widely used model format specifically among members of the COBRA modeling community [41].

MASSpy also provides support for importing and exporting models via JSON, a text-based syntax that is useful for exchanging structured data between programming languages [48]. The MASSpy JSON schema is designed for interoperability with Escher [31], a pathway visualization tool designed to visualize various -omic data sets mapped onto COBRA models. The interoperability with Escher is exploited by MASSpy to provide various pathway and node map visualization capabilities.

### Mechanistic modeling of enzyme regulation

The reconstruction of all microscopic steps performed by an enzyme (an “enzyme module”) represents the full stoichiometric description of an enzyme using mass action kinetics [22]. MASSpy facilitates the construction of enzyme modules through the EnzymeModule, EnzymeModuleForm, and EnzymeModuleReaction classes, which inherit from the MassModel, MassMetabolite, and MassReaction classes, respectively. The EnzymeModule class contains methods and attribute fields to aid in the construction of EnzymeModules based on the steps outlined for constructing enzyme modules in Du et al. [22]. Given the number and complexity of possible enzymatic mechanisms [49], MASSpy also provides the ability to group relevant objects into different user-defined categories, such as active/inactive states and different enzyme complexes. The EnzymeModuleDict objects are used to represent enzyme modules once merged into a larger model, preserving user-defined categories and other information relevant to the construction of the EnzymeModule, such as total enzyme concentration. More details can be found in the MASSpy documentation (S2 File).

### Ensemble sampling, assembly, and modeling

Ensemble approaches are used to address various issues concerning parameter uncertainty and experimental error in metabolic models [18]. Ensemble modeling refers to the assembly of dynamic models that span the feasible kinetic solution space and is useful when parameterization is incomplete or unknown, as is often the case with kinetic models. MASSpy enables ensemble modeling approaches through the use of Markov chain Monte Carlo (MCMC) sampling of fluxes and concentrations [50, 51]. The flux sampling capabilities can be derived from the COBRApy package and employ two different hit-and-run sampling methods: one with a low memory footprint [52] and another with multiprocessing support [53]. To sample metabolite concentrations, MASSpy employs a ConcSolver object to populate the optimization solver with constraints for thermodynamically feasible concentration ranges [23, 54, 55] and two hit-and-run sampling methods for concentrations were implemented in MASSpy with algorithms analogous to those for flux sampling. MASSpy provides several built-in methods for ensemble generation from sampling data. Once generated, the ensemble of models can be loaded into the MASSpy Simulation object, simulated, and visualized using built-in ensemble visualization and analysis methods. Additional details can be found in the MASSpy documentation (S2 File).

## Results

We conducted three different case studies that exemplified how MASSpy features combined to facilitate dynamic modeling of metabolism (Fig 1). In Case Study 1, we validated MASSpy as a modeling tool by describing mechanisms of enzyme regulation using enzymes modules [20]. We demonstrated the utility of the software in Case Study 2, generating an ensemble of stable kinetic models through MCMC sampling to examine biological variability while satisfying thermodynamic constraints imposed by the network. In Case Study 3, we integrated COBRA and MASS modeling methodologies to create a kinetic model of *E. coli* glycolysis from a metabolic reconstruction, providing novel insight into functional states of the proteome and activities of different isozymes. See Table 2 for a comparison of explicitly supported MASSpy features with those of other dynamic modeling tools.

**Fig 1.**
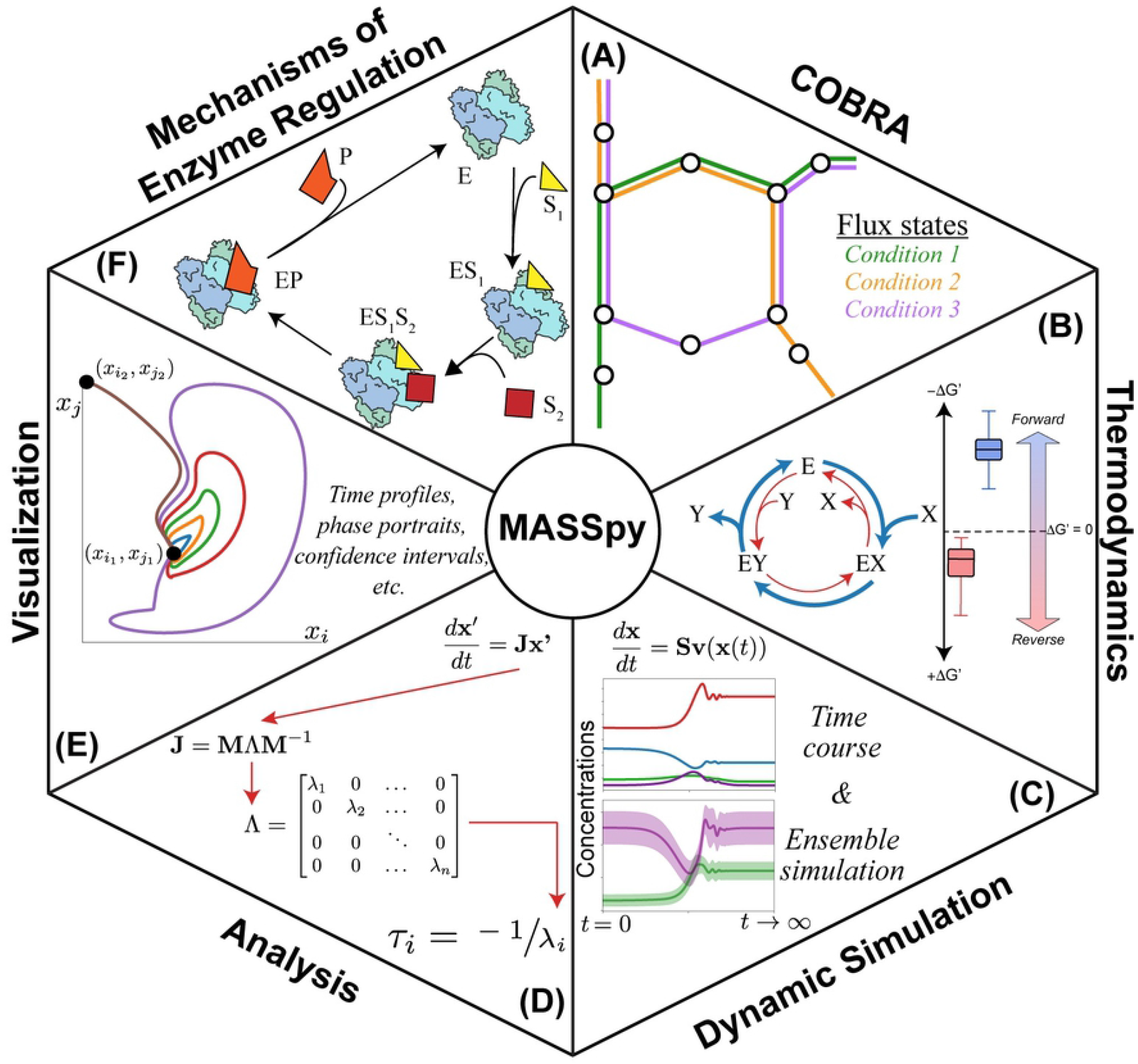
Overview of MASSpy features. (A) MASSpy expands COBRApy to provide constraint-based methods for obtaining flux states. (B) Thermodynamic principles are utilized by MASSpy to sample concentration solution spaces and to evaluate how thermodynamic driving forces shift under different metabolic conditions. (C) MASSpy enables dynamic simulation of models to characterize transient dynamic behavior and contains ensemble modeling methods to represent biological uncertainty. (D) Network properties such as relevant timescales and system stability are characterized by MASSpy using various linear algebra and analytical methods. (E) MASSpy contains built-in functions that enable the visualization of dynamic simulation results. (F) Mechanisms of enzymatic regulation are explicitly modeled in MASSpy through enzyme modules, enabling computation of catalytic activities and functional states of enzymes.

**Table 2.** Comparison of explicitly supported features for dynamic modeling tools.

| Software |  | MASSpy | MASS<br>Toolbox | Tellurium | PySB | PySCeS<br>(CMBPy) | COPASI |
| --- | --- | --- | --- | --- | --- | --- | --- |
| Version |  | 0.1.0 | 1.2.0 | 2.1.5 | 1.11.0 | 0.9.7<br>(0.7.25) | 4.27.217 |
| Environment |  | Python<br>3.6+ | Mathematica<br>9.0+ | Python<br>2.7, 3.4+ | Python<br>2.7, 3.6+ | Python<br>2.7, 3.5+ | Bindings:<br>C#, Java,<br>Python |
| Model<br>construction | Model merging | + | + | + | + |  | + |
|  | Automated rate law<br>construction | (+) only<br>mass action | (+) only<br>mass action |  | (+) only<br>mass action |  | + |
|  | Enzyme modules | + | + |  | (+)<br>monomers |  |  |
|  | Symbolic expression<br>manipulation | + | + |  |  |  |  |
|  | GPR handling | + | + | + |  | + |  |
| Sampling<br>and<br>estimation | Flux sampling | + | + |  |  |  |  |
|  | Concentration<br>sampling | + |  |  |  |  |  |
|  | Parameter<br>estimation | (+) PERCs | (+) PERCs |  |  |  | + |
| Simulation | ODE | + | + | + | + | + | + |
|  | Stochastic |  |  | + | + |  | + |
| Analysis | Steady state | + | + | + | (+) via<br>simulation | + | + |
|  | Stoichiometric | + | + | + |  | + | + |
|  | Sensitivity |  |  | + | + | + | + |
|  | Metabolic control |  |  | + |  | + | + |
|  | Stability | + | + | + |  | + | + |
|  | Gene knockouts | + | + |  |  | + |  |
|  | Flux balance / flux<br>variability | + | + |  |  | + |  |
| Visualization | Time profiles | + | + | + |  | + | + |
|  | Phase portraits | + | + | + |  | + | + |
|  | Pathway maps | (+) via<br>Escher | + | (+) via<br>Graphviz | (+) via<br>Graphviz | (+) via<br>FAME | + |
| Quality<br>control and<br>assurance | Elemental balancing | + | + |  |  | + |  |
|  | Thermodynamic<br>feasibility | + | + |  |  |  |  |
|  | Assistance w/<br>undefined values | + | + |  |  |  |  |
| Standards | SBML | + | + | + | + | + | + |
|  | SED-ML |  |  | + |  | (+) export | (+) export |
|  | COMBINE |  |  | + |  | (+) export | + |

“+” denotes explicit support for the feature. “(+)” denotes explicit support via an interface, or with some limitations. Explicit support is determined based on whether clear methods and documentation demonstrating feature support are available from the listed software. For example, although MASSpy could utilize the stochastic simulation capabilities of libRoadRunner, no MASSpy methods or documentation providing explicit support for stochastic simulation in MASSpy exist currently.

### Case Study 1: Enzyme regulation in MASS models

Here, we demonstrated MASSpy as a modeling tool and the MASSpy implementation of enzyme modules by replicating the results produced by Yurkovich et al. [20]. The authors used the MASS Toolbox [25] to elucidate the systems-level effects of allosteric regulation. We used MASSpy to reconstruct enzyme modules for hexokinase, phosphofructokinase, and pyruvate kinase in RBC glycolysis using the same mechanisms as previously described [20]. We provided several in-depth tutorials for constructing MASS models and enzyme modules in MASSpy, which can be found in the documentation (S2 File).

We integrated the reconstructed enzyme modules into the glycolytic model to introduce varying levels of regulation. Because enzyme modules were constructed and parameterized for the steady-state conditions of the MASS model, addition of an enzyme module to a MASS model was a straightforward and scalable process. The overall reaction representation for the enzyme in the MASS model was removed and replaced with the set of reactions that comprise the microscopic steps of the enzyme module (Fig 2) We performed dynamic simulations, subject to physiologically relevant perturbations, to provide a fine-grained view of the concentration and flux solution profiles for individual enzyme signals and qualitatively represent the systemic effects of additional regulatory mechanisms. We then used MASSpy visualization methods to replicate key results [20] (S3 File). Through this case study, we have demonstrated how enzyme modules were constructed from enzymatic mechanisms in MASSpy, and we validated MASSpy as a dynamic modeling tool by exploring previously reported systems-level effects of regulation [20]. See S3 File for all data and scripts associated with this case study, including kinetic parameters for all three enzyme modules.

**Fig 2.**
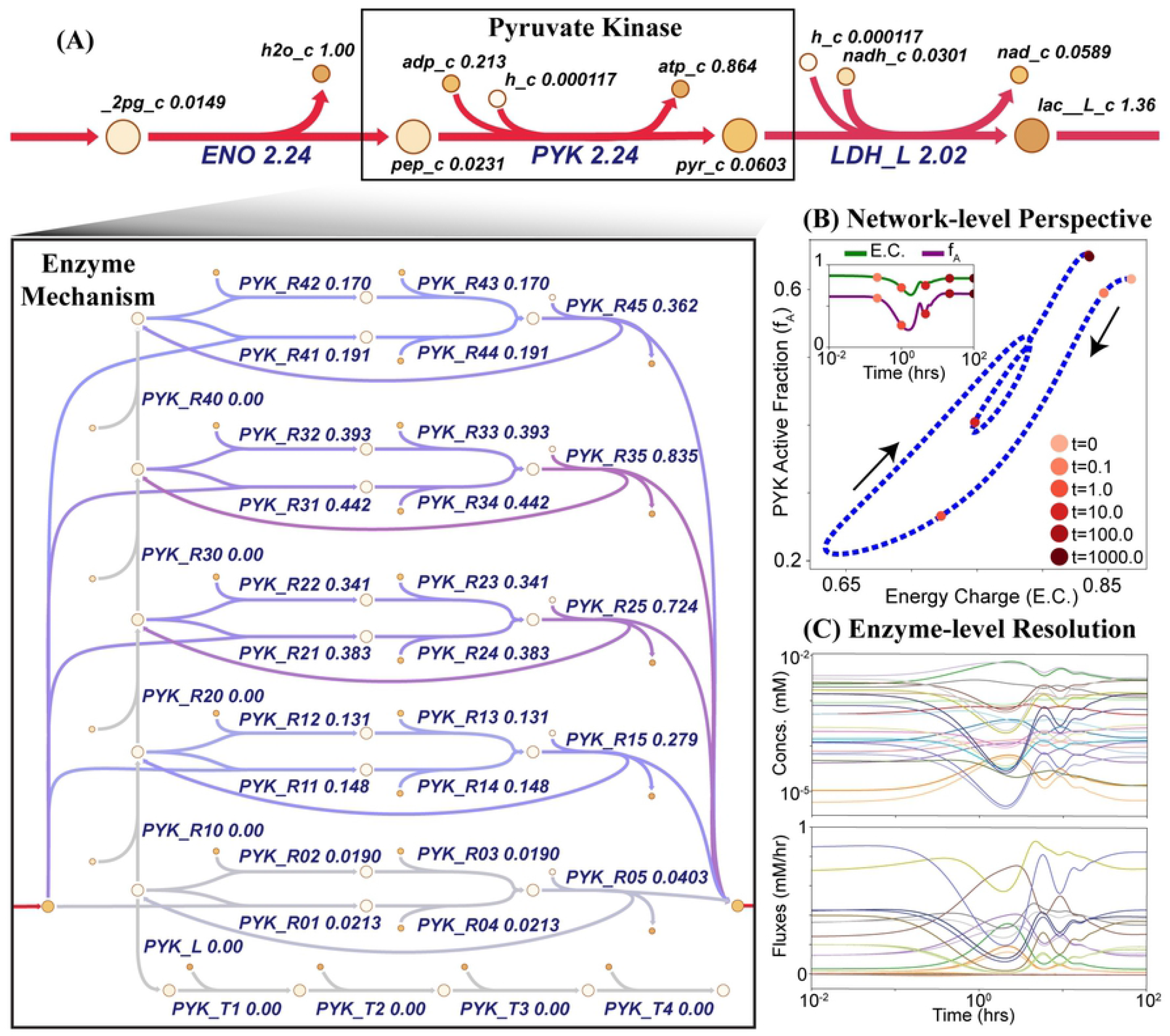
Enzyme modules are explicit representations of enzymatic regulatory mechanisms. (A) The reaction catalyzed by pyruvate kinase is replaced with the stoichiometric description of the enzymatic mechanism. The steady state values obtained after simulating a 50% increase of ATP utilization are mapped onto a metabolic pathway map drawn using Escher [31]. The colors represent flux values and range from red to purple to gray, with red indicating higher flux values and gray indicating lower flux values. (B) Enzyme modules provide a network-level perspective of regulation mechanisms by plotting systemic quantities against fractional states of enzymes as described in Yurkovich et al. [20]. (C) The different signals of the enzyme module can be observed to provide enzyme-level resolution of the regulatory response.

### Case Study 2: Ensemble sampling, assembly, and modeling

Many ensemble modeling approaches utilize sampling methods to approximate missing values and quantify uncertainty in metabolic models [17, 18, 23, 51, 56, 57]. To demonstrate the sampling and ensemble handling capabilities of MASSpy, we utilized MCMC sampling with an ensemble modeling approach to assess the dynamics for a range of pyruvate kinase enzyme modules (Fig 3). Using RBC glycolysis and hemoglobin as the reference model [20] we used MCMC sampling to generate 25 candidate flux states and 25 candidate thermodynamically feasible concentration states, allowing for variables to deviate from their reference state by up to 80 percent. This procedure resulted in 625 models that represent all possible combinations of flux and concentration states. We calculated pseudo-elementary rate constants (PERCs) and steady-state for each model. We simulated a 50% increase in ATP utilization to mimic a physiologically relevant disturbance, such as increased shear stress due to arterial constriction [58]. Out of 625 models, 15 were discarded due to an inability to reach a steady state.

**Fig 3.**
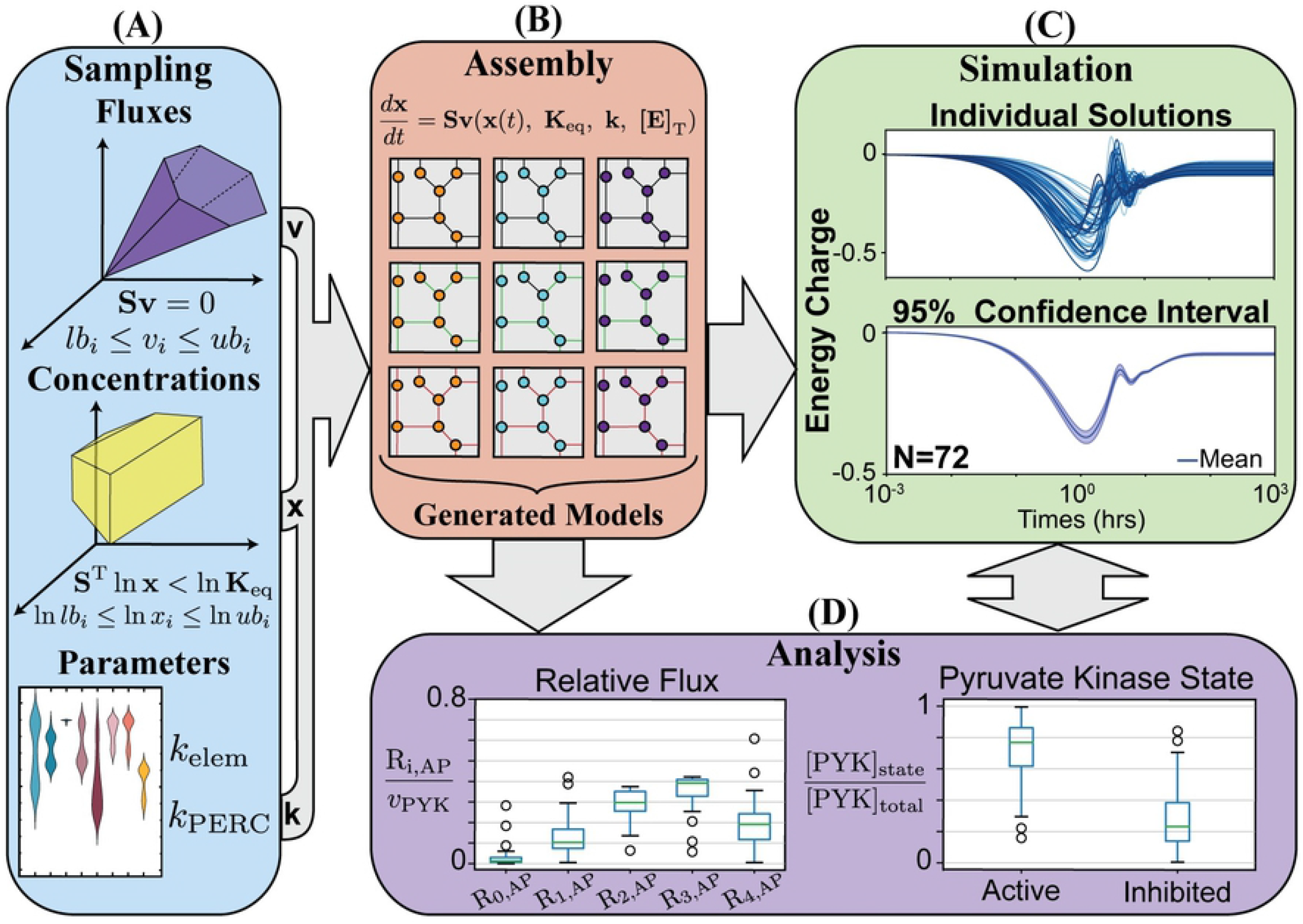
A workflow for ensemble creation and modeling using MCMC sampling. The general process for generating and assembling an ensemble of models for dynamic simulation and analysis. (A) The solution spaces for fluxes and concentrations are sampled using MCMC sampling to generate data for ensemble creation. Rate constants are obtained through parameter fitting for elementary rate constants and computation of PERCs in addition to MCMC sampling. (B) Sampling data is integrated into models, producing an assembly of models with variations in flux and concentration states. After models are created, ensembles of models can be studied through (C) dynamic simulation and (D) analysis.

We then reconstructed enzyme modules for pyruvate kinase [20] for the remaining 610 models. Numerical values of rate constants for each enzyme were determined using the SciPy implementation of a trust-region method for nonlinear convex optimization [59]. Without knowledge of physiological constraints, the numerical solutions for rate constants produced by optimization routine were not guaranteed to be physiologically feasible. Therefore, we integrated these enzyme modules into their MASS models and simulated with and without the ATP utilization increase to filter out infeasible models based on whether they could reach a stable steady state. Out of 610 models, 446 could not reach a steady state and 92 could not reach a steady state with the perturbation, leaving 72 stable models for ensemble simulation and analysis.

The time-course results for the ensemble energy charge deviation were plotted with a 95% confidence interval. From these results, it can be seen that the mean energy charge decreased at most about 40% from its baseline value (Fig 3C). Steady state analysis of the pyruvate kinase enzyme modules after the disturbance revealed a strong preference for the enzyme to remain in an active state, with a median value of approximately 77%. The differences in candidate flux and concentration states resulted in an interquartile range between 62-86% of total pyruvate kinase, with nearly all variations of pyruvate kinase in the ensemble maintaining a steady state value of at least 30% active. Furthermore, examination of the relative flux load through the R_i,AP_ forms showed that most of the flux load was carried by the R_2,AP_ and R_3,AP_ reaction steps while a miniscule fraction was carried by the R_0,AP_ reaction step, regardless of variation. However, the variations had an effect on whether R_2,AP_ carried more flux than R_3,AP_, and whether the remaining flux was predominantly distributed through the R_1,AP_ or the R_4,AP_ reaction step. Through this case study, we have demonstrated how MASSpy sampling facilitated the assembly and simulation of an ensemble to characterize the dynamic response of a key regulatory enzyme and quantify its functional states after a physiologically relevant disturbance. See S3 File for all data and scripts associated with this case study.

### Case Study 3: Computing functional states of the *E. coli* proteome

Here, we illustrated unique features of MASSpy in a workflow to compute the functional states of the proteome, providing insight into distribution of catalytic activities of enzymes for different metabolic states. We utilized COBRA and MASS modeling methodologies to incorporate omics data into a metabolic reconstruction of *E. coli*, formulating a kinetic model containing all microscopic steps for each enzymatic reaction mechanism of the glycolytic subnetwork. Once formulated, we interrogated the model to examine the shift in thermodynamic driving force for *E. coli* on different carbon sources and to compare the activities of different isozymes, exemplifying the utility of MASSpy.

To construct a kinetic model of the glycolytic subnetwork, we integrated steady-state data for growth on glucose and pyruvate carbon sources from Luca et. al [60] into the iML1515 genome-scale metabolic reconstruction of *E. coli* K-12 MG1655 [61]. For each carbon source, we fixed the growth rate for iML1515 and performed FBA using a quadratic programming objective to compute a flux state that minimized the error between known and computed fluxes. For the irreversible enzyme pairs of phosphofructokinase/fructose 1,6-bisphosphatase (PFK/FBP) and pyruvate kinase/phosphoenolpyruvate synthase (PYK/PPS), individual flux measurements were each increased by 10% of the net flux for the enzyme pair without changing the overall net flux value to ensure presence of the enzyme as seen in proteomic data [62].

Once the flux state was obtained for each carbon source, the glycolytic subnetwork was extracted from iML1515. Flux units were converted into molar units using volumetric measurements of *E. coli* obtained from Volkmer and Heinemann [63], and equilibrium constants obtained from eQuilibrator [63] through component contribution [64] were set for each reaction. Concentration growth data from Luca et. al [60] was integrated into the model and minimally adjusted for thermodynamic feasibility; for metabolites missing concentration data, an initial value of 0.001 M was provided before adjustments through sampling. Concentrations were sampled within an order of magnitude of their current value to produce an ensemble of 100 candidate models for each growth condition. Metabolite sinks were added to the model to account for metabolite exchanges between the modeled subnetwork and the global metabolic network outside of the scope of the model.

For each model in the ensemble, enzyme modules were constructed for each reaction using a nonlinear parameter fitting package (https://github.com/opencobra/MASSef) and kinetic data extracted from the literature. Additional isozymes of phosphofructokinase (PFK), fructose 1,6-bisphosphatase (FBP), fructose 1,6-bisphosphate aldolase (FBA), phosphoglycerate mutase (PGM), and pyruvate kinase (PYK) were also constructed, bringing the total amount of enzyme modules to 17. Fluxes through individual isozymes were set by splitting the steady state flux between the major and minor isozyme forms. After integrating all enzyme modules into the network, each model was simulated to steady state and exported for analysis.

The Gibbs free energy for each enzyme-catalyzed reaction and fractional abundance of each enzyme form were calculated. Sensitivity analysis of the flux split between the isozyme forms revealed that the Gibbs free energy and fractional abundance of enzyme forms did not show significant variation for either carbon source (S1 Fig); therefore, remaining analyses were done with 75% and 25% of the flux assigned to major and minor isozyme forms, respectively.

Comparison of the glucose and pyruvate growth conditions revealed that the free energy of the reversible reactions remained close to equilibrium, changing from one metabolic state to another as the thermodynamic driving force shifts according to the carbon source, as seen in reversible reactions phosphoglucoisomerase (PGI), triose phosphate isomerase (TPI), glucose 6-phosphate dehydrogenase (GAPD), phosphoglycerate kinase (PGK), phosphoglycerate mutase (PGM), and enolase (ENO). (Fig 4A) The reaction pairs, PFK/FBP and PYK/PPS, maintained their flux directions to form a futile cycle across conditions

**Fig 4.**
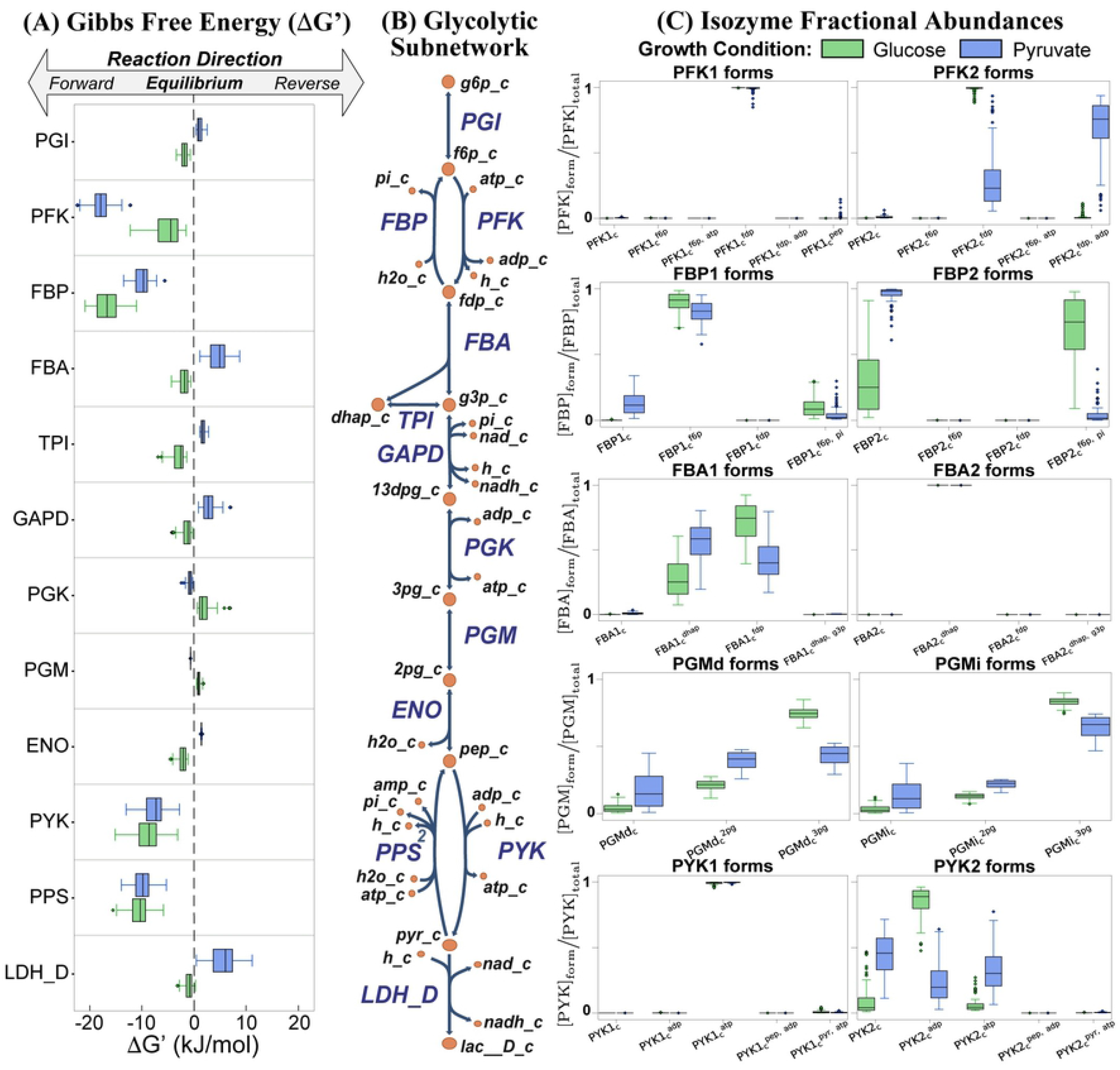
Comparison of free energy and isozyme fractional abundances for carbon sources. (A) The Gibbs free energy represents the thermodynamic driving force, shifting the metabolic state depending on the carbon source. (B) The glycolytic subnetwork extracted from *E. coli* iML1515 consists of 12 reactions represented by the 17 enzyme modules. (C) The fractional abundance for each enzyme form can be computed and compared for the different isozyme pairs, providing insight into how the catalytic activity is distributed across the isozymes in glucose and pyruvate growth conditions. The fractional abundances for all enzymes can be found in the supplement (S2 Fig)

Steady state analysis of the isozyme fractional abundances elucidated a preference for a specific enzyme state conserved among growth conditions for PFK1, FBA2, and PYK1 (Fig 4C). Steady state analysis also revealed that the isozyme pairs of PFK, FBP, and PYK primarily existed in their product-bound form, a reflection of the metabolite concentration levels found in S4 File. Specifically, the relatively high concentrations of fructose 1,6-diphosphate (FDP) observed for glucose growth conditions and of adenosine triphosphate (ATP) observed for pyruvate growth conditions contributed to significant differences between enzyme product and reactant concentration levels, creating the conditions favorable to the product-bound enzyme forms. Both the major and minor PGM isozymes have a similar distribution in their enzyme forms. The fractional abundance of all enzyme states in the glycolytic subnetwork can be found in S2 Fig. Through this case study, we have demonstrated that MASSpy can be used to gain insight into the distribution of functional states for the glycolytic proteome in *E. coli* without prior knowledge of enzyme concentrations. See S3 File for all data and scripts associated with this case study, including microscopic steps and kinetic parameters for all enzyme modules

## Discussion

We describe MASSpy, a free and open-source software implementation for dynamic modeling of biological systems. MASSpy expands the COBRApy framework, leveraging existing methods familiar to COBRA users combined with kinetic modeling methods to form a single, intuitive framework for constructing and interrogating dynamic models. In addition to enabling dynamic simulation, MASSpy contains tools for facilitating the reconstruction and analysis of enzyme modules, MCMC sampling and ensemble modeling capabilities, interfacing with packages for pathway visualization (Escher, [31]), and exchanging models in SBML format (libSBML [33]). Taken together, the presentation of the MASSpy software package has several important implications for practitioners of dynamic metabolic simulation.

MASSpy provides several benefits over existing modeling packages (Table 2). While MASS models provide an algorithmic approach for generating dynamic models that has already proven useful in several metabolic studies [20–24], a formal implementation of the MASS framework has only existed on a commercial software platform (Mathematica). MASSpy’s seamless integration with COBRApy offers a vast array of constraint-based and dynamic modeling tools within a single open-source framework. MASSpy primarily utilizes the MASS approach and therefore integrates a suite of tools into its framework for addressing issues specific to MASS modeling. Unlike other packages for traditional kinetic modeling, MASSpy incorporates both COBRA methods and MCMC sampling methods for estimating missing values for several data types. Furthermore, MASSpy contains unique capabilities to facilitate the construction and analysis of detailed enzyme modules (i.e., microscopic steps), which allow for the dynamics of transient responses to be observed in situations in which the quasi-steady state and quasi-equilibrium assumptions cannot be applied. By directly expanding the COBRApy framework for MASSpy, current COBRApy users will find that MASSpy provides procedures and protocols that they may be familiar with, and allows members of the COBRA community to directly integrate new tools into their existing workflows.

MASSpy is primarily built for deterministic simulations of a metabolic model and thus may face limitations for other uses. For example, a package like PySCeS [65] could be utilized to perform stochastic simulations. Users who often analyze sensitivity may prefer Tellurium and its implementation of libRoadRunner [32, 66] for explicit support of metabolic control analysis (MCA) workflows; however, MASSpy does contain similar MCA methods through its own implementation of libRoadRunner. Other dynamic modeling packages offer certain features not available in MASSpy, such as a graphical user interface (COPASI [67]) or a rule-based modeling approach (PySB [29]): see Table 2 for a comparison of MASSpy’s software features with other existing dynamic modeling packages. MASSpy’s use of SBML facilitates the transfer of models to other software environments if desired [45].

Taken together, we have described MASSpy, a Python-based software package for the reconstruction, simulation, and visualization of dynamic metabolic models. MASSpy provides a suite of dynamic modeling tools while leveraging existing implementations of constraint-based modeling tools within a single, unified framework. The case studies presented here validate MASSpy as a modeling tool and demonstrate how the combination of constraint-based and kinetic modeling features support data-driven solutions for various dynamic modeling applications. We anticipate that the community will find MASSpy to be a useful tool for dynamic modeling of metabolism.

## Availability and future directions

### Software availability and requirements

MASSpy version 0.1.0 is available as a Python package hosted on the Python Package Index (https://pypi.org/project/masspy/), licensed under GNU General Public License, version 3.0 (GPL-3.0). All external dependencies integrated and utilized by MASSpy are also available on the Python Package Index (https://pypi.org/) and are licensed under their respective licensing terms. Both the Gurobi Optimizer (Gurobi Optimization, Houston, TX) and the CPLEX Optimizer (IBM, Armonk, NY) are freely available for academic use, with solvers and installation instructions found at their respective websites. The latest source code for MASSpy is currently available on GitHub (https://github.com/SBRG/MASSpy) and is compatible with Mac OS X, Linux, and Windows operating systems. Instructions for MASSpy installation can be found in the repository README or in the documentation (S2 File). The data, scripts, and instructions needed to reproduce the results of the case studies can be found in the S3 File.

### Documentation

The documentation for MASSpy is available online (https://masspy.readthedocs.io/). Good documentation is vital to the adoption and success of a software package; it should teach new users how to get started while showing more experienced users how to fully capitalize on the software’s features [41, 68]. For new users, MASSpy provides several simple tutorials demonstrating the usage of MASSpy’s features and its capabilities. The MASSpy documentation also contains a growing collection of examples that demonstrate the use of MASSpy, including examples of workflows, advanced visualization tutorials, and in-depth textbook [24] examples that teach the fundamentals for dynamic modeling of mass action kinetics (S2 File).

### Improvements and community outreach

The MASSpy package is designed to provide various dynamic modeling tools for the openCOBRA community; therefore a substantial portion of future development for MASSpy will be tailored toward fulfilling the needs of the COBRA community based on user feedback and feature requests. New MASSpy releases will utilize GitHub for version control and adhere to Semantic Versioning guidelines (https://semver.org/) in order to inform the community about the compatibility and scope of improvements. Examples of potential improvements for future releases of MASSpy include bug fixes, additional SBML compatibility, new import/export formats, support for additional modeling standards, explicit support for additional libRoadRunner simulation capabilities, and implementation of additional algorithms relevant to MASS modeling approaches. As the systems biology field continues to address challenges in dynamic models of metabolism, MASSpy will continue to expand its collection of modeling tools to support data-driven reconstruction and analysis of mechanistic models.

## Supporting information

**S1 Fig. Sensitivity analysis on flux split through isozymes in the** *E. coli* **glycolytic subnetwork**. The Gibbs free energy of enzyme-catalyzed reactions and the fractional abundance for isozyme states for all isozyme pairs for (A) glucose growth conditions and (B) pyruvate growth conditions when computing the functional states of the *E. coli* proteome in Case study 3.

**S2 Fig. Fractional abundance of all enzyme states in the** *E. coli* **glycolytic subnetwork**. The fractional abundance for all enzymes states of all enzyme modules when computing the functional states of the *E. coli* proteome in Case Study 3.

**S1 File. The source code for MASSpy version 0.1.0**. The latest version of the MASSpy software can be found at https://github.com/SBRG/MASSpy. (ZIP)

**S2 File. The documentation for MASSpy version 0.1.0** The latest version of the MASSpy documentation can be found at https://masspy.readthedocs.io. (ZIP).

**S3 File. Data and Jupyter notebooks for case studies**. All files necessary to repeat each case study. Each folder contains the relevant data, scripts, and Jupyter notebooks for that case study. Alternatively, these files can be found at https://github.com/SBRG/MASSpy-publication (ZIP)

**S4 File. Steady state concentration data in the** *E. coli* **glycolytic subnetwork**. Includes the steady state concentration data for all metabolites and enzymes in the *E. coli* glycolytic subnetwork in Case Study 3. (XLSX)

## Acknowledgments

The authors gratefully acknowledge: Patrick Phaneuf and Laurence Yang for discussions about software design considerations during the development of the MASSpy software, Zak King for help concerning Escher interoperability and general software development, Bin Du for discussions about the implementation of enzyme modules and thermodynamic feasibility features, Colton Lloyd for discussions about COBRA methods and expanding the COBRApy framework, and the users who provided feedback during the development process for the initial MASSpy release.

